# Aneurysmal Growth in Type-B Aortic Dissection: Assessing the Impact of Patient-Specific Inlet Conditions on Key Haemodynamic Indices

**DOI:** 10.1101/2023.02.12.528194

**Authors:** C. Stokes, D. Ahmed, N. Lind, F. Haupt, D. Becker, J. Hamilton, V. Muthurangu, H. von Tengg-Kobligk, G. Papadakis, S. Balabani, V. Díaz-Zuccarini

## Abstract

Type-B Aortic Dissection is a cardiovascular disease in which a tear develops in the intimal layer of the descending aorta, allowing pressurized blood to delaminate the layers of the vessel wall. In medically managed patients, long-term aneurysmal dilatation of the false lumen (FL) is considered virtually inevitable and is associated with poorer disease outcomes. While the pathophysiological mechanisms driving FL dilatation are not yet understood, hemodynamic factors are believed to play a key role. Computational Fluid Dynamics (CFD) and 4D-Flow MRI (4DMR) analyses have revealed correlations between flow helicity, oscillatory wall shear stress, and aneurysmal dilatation of the FL. In this study, we compare CFD simulations using a patient-specific, three-dimensional, three-component inlet velocity profile (3D IVP) extracted from 4DMR data against simulations with flow rate-matched uniform and axial velocity profiles that remain widely used in the absence of 4DMR. We also evaluate the influence of measurement errors in 4DMR data by scaling the 3D IVP to the degree of imaging error detected in prior studies. We observe that oscillatory shear and helicity are highly sensitive to inlet velocity distribution and flow volume throughout the FL and conclude that the choice of IVP may greatly affect the future clinical value of simulations.

## 1 Background

Type-B Aortic Dissection (TBAD) is treated medically in the absence of complications. Despite the use of anti-hypertensive treatment, aneurysmal growth is observed in up to 87% of these medically treated patients^1^, usually with more pronounced growth in the thoracic aorta^2^. Aortic growth is a known risk factor for late adverse events including aortic rupture, so there is value from a clinical perspective in understanding how and why this growth occurs; predictors of aneurysmal growth may be used to tailor treatment and improve disease outcomes^1^. Unfortunately, the pathological mechanisms driving aneurysmal growth are not yet well-understood and existing anatomical predictors, including a large false lumen (FL) diameter at presentation and a large, proximally located primary entry tear (PET)^3^, perform poorly^4^; better clinical risk stratification tools are needed.

Haemodynamic quantities such as pressure and wall shear stress (WSS) may offer greater predictive power than anatomical markers as they are more directly linked with the physiological mechanisms at play; WSS characteristics are known to influence the behaviour of endothelial cells while fluid pressure can affect the regulation of arterial structure and inflict further delamination of the aortic wall in TBAD^5^. Greater FL flow^6^ and pressurisation^7^ have been associated with aortic growth in addition to retrograde flow through the PET^4^ which has been associated with increased FL pressurisation. Regions of high oscillatory shear index (OSI) and low time-averaged WSS (TAWSS) have been linked with aneurysmal growth and rupture in TBAD^6^, creating a feedback loop of degradation as growth further exacerbates these effects^8^.

Despite the growing evidence of haemodynamic involvement in TBAD, links between flow and disease progression remain inconsistent and robust, large-scale clinical trials are needed to develop clinically applicable haemodynamic predictors. 4D-Flow MRI (4DMR) and Computational Fluid Dynamics (CFD) are the two predominant means of haemodynamic analysis, but further efforts to understand and limit the errors and uncertainties associated with these modalities are required. 4DMR is known to perform poorly in low-velocity regions^9^ while high velocity gradients cannot be adequately captured due to low spatiotemporal resolution, leading to significant errors in extracted WSS indices^10^ and limiting its use in investigating flow-mediated vascular remodelling. CFD offers high spatiotemporal resolution but patient-specific accuracy is strongly influenced by segmentation quality, the choice of boundary conditions and numerous other modelling assumptions. Applying patient-specific boundary conditions in CFD from 4DMR data is currently the most favourable means of producing accurate, high-fidelity haemodynamic data. As 4DMR data is not routinely acquired, simplified or literature-based boundary conditions are frequently used, despite the knowledge that this can have a profound impact on the accuracy of the results.

While the impact of inlet conditions on velocity magnitude and TAWSS have been investigated in TBAD^11,12^, these efforts did not consider oscillatory shear indices or flow helicity, each of which is associated with aneurysmal growth^6,13^ and has shown substantial sensitivity to inlet conditions in healthy aortae^14^. In this study, we explore the impact of several widely used inlet conditions on disturbed shear indices and flow helicity in a case of chronic TBAD. In this patient, widespread aneurysmal growth of up to 88% was observed in the FL over a two-year period. We compare the gold-standard three-dimensional, three-component inlet velocity profile (3D IVP) extracted from 4DMR data with flow rate-matched flat (F) and through-plane (TP) profiles that remain widely used in the absence of 4DMR. Furthermore, we assess the impact of 4DMR imaging errors on the bulk flow solution by modulating the measured inlet velocity components by the degree of velocity underestimation observed in previous studies ^15^.

To further examine the impact of inlet conditions on the velocity field, we employ Proper Orthogonal Decomposition (POD), a reduced-order modelling (ROM) technique that is becoming more widely used to analyse cardiovascular flows ^16^. POD is typically applied to identify coherent flow structures which optimally capture the fluctuating kinetic energy (KE) of the velocity field, thus facilitating a deeper understanding of complex flows ^17^. POD has been applied to characterise turbulence in cerebral arteries^18^, examine the impact of inflow strength and angle in cerebral and abdominal aortic aneurysms and identify differences between healthy and pathological flow conditions within them^16,19^. It has also been used to enhance the resolution of 4DMR^20^.

## 2 Methods

### 2.1 Clinical data

Medical imaging data from a 56-year-old male patient with chronic TBAD were acquired at [REDACTED UNTIL ACCEPTANCE] under ethical approval from the local Institutional Review Board (ID: 2019-00556). A Siemens SOMATOM Definition Flash (Siemens AG, Munich, Germany) was used to acquire Computed Tomography Angiography (CTA) data with an isotropic spatial resolution of 0.5 mm. Four months after CTA, 4D-Flow MRI (4DMR) data were acquired using a Siemens Aera 1.5T, and again at two years after CTA, using a spatial resolution of 2.25 x 2.25 x 3.00 mm, a velocity encoding (VENC) of 150 cm/s and 16 timeframes across the cardiac cycle. A heart rate of 94 beats per minute was extracted from the 4DMR data and a single brachial measurement of 138/81 Pa was obtained before the first 4DMR acquisition. Luminal area measurements were extracted from each 4DMR dataset at 5mm increments along the thoracic aorta, indicating that over the two years between acquisitions, the thoracic FL dilated by an average of 35% and a maximum of 88%.

### 2.2 Segmentation & Meshing

The computational domain, extending from the ascending aorta to the distal end of the dissection at the external and internal iliac arteries, was manually segmented from CTA data using Simpleware ScanIP (Synopsys Inc., CA, USA) and Autodesk Meshmixer (Autodesk Inc., CA, USA) and non-rigidly registered to the 4DMR domain using a continuous-point-drift algorithm in MATLAB (Matlab, Natick, MA, USA)^21^. For later analysis, the volume was divided into five sections: the ascending aorta (AA), the thoracic TL and FL (TL*_t_* and FL*_t_*), and the abdominal TL and FL (TL*_a_* and FL*_a_*) as shown in Fig. 1.

**Figure 1:**
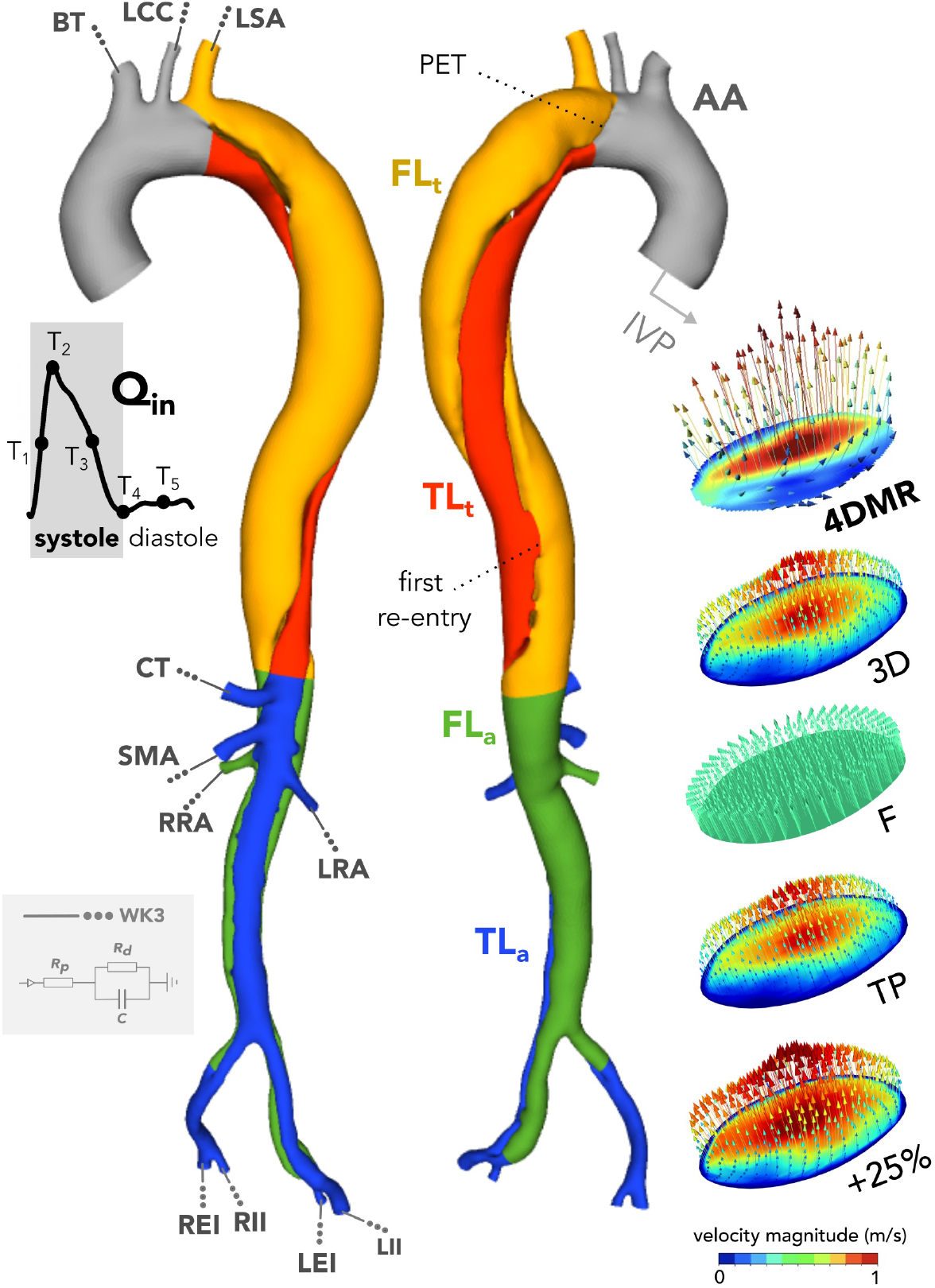
The computational domain with schematic depictions of inlet and outlet boundary conditions. The thoracic and abdominal regions of the TL and FL are coloured separately and denoted by subscripts a and t, respectively. The velocity magnitude contours and vectors at peak systole of each patient-specific inlet velocity profile are provided bottom right. The coloured regions of the aortic geometry represent the analysis regions used for helicity computations.

The volume was meshed using tetrahedral elements in Fluent Mesh (ANSYS Inc., PA, USA). Mesh sizing parameters were determined with a mesh independence study described in Supplementary Material SM1. The final mesh contained 2.30M elements, including ten near-wall layers and a first-cell height corresponding to a mean y+ of 0.83 at peak systole.

The primary entry tear (PET) is located at the left subclavian artery (LSA) and is 18 mm in diameter. The first and largest re-entry tear is located 170 mm distal to the PET, as shown in Fig. 1. Nineteen further luminal communications are present.

### 2.3 Inlet conditions

We will compare simulations with four commonly used patient-specific inlet velocity profiles (IVP), each derived from the first: a three-component, three-dimensional (3D) IVP which precisely matches the magnitude and direction of 4DMR velocities at the inlet plane, as shown in Fig. 1. A flat (F) IVP was generated with an identical inlet flow waveform (*Q_in_*), but with a uniform velocity distribution in the plane-normal direction, as is typically deployed when flow MRI data is not available. Next, an axial or through-plane (TP) IVP was produced, comprising only the plane-normal component of the 3D IVP such that *Q_in_* was matched. TP IVPs may be used when only 2D-Flow MRI data is available. The final profile was identical to the 3D IVP, but with each velocity component increased by 25%; 4DMR data has been shown to underestimate peak flow rate and velocity by *≈*20-30%^15^ depending on image resolution. The stroke volume of this IVP is thus 25% higher than the other three IVPs.

To produce the 3D IVP, velocity data from 4DMR was extracted at the inlet plane using GTFlow (Gyrotools LLC, Zurich, CH). Individual contours were manually generated at each imaging time point to track the ascending aortic wall as it translated and expanded across the cardiac cycle. As the CFD inlet shape and location are fixed in time, and simulations require a higher temporal and spatial resolution than 4DMR provides, an algorithm was developed to register and interpolate the 4DMR data onto the CFD inlet to generate the 3D IVP. First, the 4DMR contour region from each timeframe was mapped to the CFD inlet using non-rigid continuous point drift in MATLAB^21^. By definition, non-rigid registration does not preserve the distance between spatial points and may distort the velocity profile. To minimise the impact of this effect, rigidity parameters were increased. Furthermore, 4DMR contours regions closely matched the circular shape of the CFD inlet, thus preventing excessive distortion. Each registered component velocity field at each timestep was then spatially interpolated onto a fixed, uniform grid to facilitate temporal interpolation to match the simulation timestep. Temporal spline interpolation was performed over numerous cardiac cycles to ensure a smooth flow waveform. Points at the perimeter of the inlet were set to zero to ensure that the non-slip wall boundary condition was met throughout the domain. Finally, spatial interpolation of the IVP onto the inlet mesh was performed automatically by the solver, ANSYS CFX 2020.

### 2.4 Outlet boundary conditions

Patient-specific flow and pressure distributions were reproduced in each simulation using three-element Windkessel (WK3) outlet boundary conditions. Target systolic and diastolic pressures (*P_s_* and *P_d_*) were derived as follows:

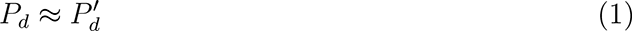

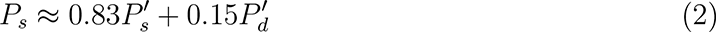

where 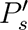 and 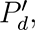 are the patient’s brachial systolic and diastolic pressure measurements^22^. The same pressure targets were used for all simulations.

Mean outlet flow rates were extracted from 4DMR at each major branch using GTFlow. Due to lower relative image resolution, measurement uncertainties in 4DMR are higher in these smaller branches than the aorta. To mitigate the effects of any associated imaging errors, the mean flow rate at each outlet was normalised by the mean flow difference between aortic planes upstream and downstream of its containing group of branches (supra-aortic, abdominal, iliac). For the +25% case, target 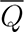 was increased by 25% at each outlet.

Using these pressure and flow targets and the inlet waveform extracted from the 3D IVP, WK3 parameters were tuned using a 0D lumped parameter model of the aorta in 20-sim (Controllab, Enschede, NL) using our previously-developed technique^23^. The final WK3 parameters are shown in Table 1 alongside the target and simulated mean outlet flow rates for each IVP 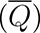. WK3 parameters were identical in the 3D, F and TP cases, but adjusted to maintain the target pressure range in the +25% case.

**Table 1:** Outlet mean flow rates and WK3 parameters in each simulation and at each outlet shown schematically in Fig. 1. ρ indicates the proportion of R_tot_ attributed to R_1_ in each WK3. WK3 parameters were identical between 3D, F and TP, but altered in +25% to account for the additional inlet flow volume.

| | | $\bar{Q}$<br>ml/s | | | | | $R_{WK3}^{tot}$<br>mmHg/ml/s | | $C_{WK3}$<br>ml/mmHg | |
| --- | --- | --- | --- | --- | --- | --- | --- | --- | --- | --- |
| | Target | 3D | F | TP | +25% | $\rho$ | 3D/F/TP | +25% | 3D/F/TP | +25% |
| BT | 17.10 | 17.69 | 17.00 | 17.47 | 22.11 | 0.030 | 5.77 | 4.37 | 0.24 | 0.63 |
| LCC | 3.84 | 3.97 | 3.77 | 3.89 | 4.88 | 0.030 | 25.67 | 19.43 | 0.05 | 0.14 |
| LSA | 6.69 | 6.98 | 6.63 | 6.83 | 8.57 | 0.030 | 14.75 | 11.16 | 0.09 | 0.25 |
| CT | 16.31 | 16.14 | 15.56 | 16.06 | 19.95 | 0.056 | 6.05 | 4.58 | 0.22 | 0.59 |
| SMA | 14.09 | 13.86 | 13.41 | 13.85 | 17.24 | 0.056 | 7.00 | 5.30 | 0.19 | 0.51 |
| LR | 16.59 | 15.63 | 15.49 | 15.72 | 19.00 | 0.280 | 5.94 | 4.50 | 0.23 | 0.61 |
| RR | 12.64 | 12.25 | 12.02 | 12.19 | 14.88 | 0.280 | 7.80 | 5.90 | 0.17 | 0.46 |
| LEI | 9.61 | 9.60 | 9.26 | 9.56 | 11.85 | 0.056 | 10.26 | 7.76 | 0.13 | 0.35 |
| LII | 4.76 | 4.76 | 4.58 | 4.72 | 5.90 | 0.056 | 20.72 | 15.68 | 0.07 | 0.17 |
| REI | 5.21 | 5.41 | 5.15 | 5.29 | 6.69 | 0.056 | 18.95 | 14.34 | 0.07 | 0.19 |
| RII | 3.01 | 3.17 | 3.00 | 3.07 | 3.98 | 0.056 | 32.79 | 24.81 | 0.04 | 0.11 |

### 2.5 Simulation

Transient simulations were performed with ANSYS CFX 2020 R2 using timesteps of 1 ms until cyclic periodicity was reached, which we defined as *<* 1% change in systolic and diastolic pressure between subsequent cycles. The Reynolds-Averaged Navier-Stokes and continuity equations were solved numerically using the implicit, second-order backward-Euler method and a root-mean-square residual target of 10*^−^*^5^ for all equations within each timestep. Walls were modelled as rigid with a no-slip condition as Cine-MRI data was not available to tune patient-specific aortic compliance using our previously developed Moving Boundary Method^24,23^.

Blood was modelled as an incompressible, non-Newtonian fluid using the Tomaiuolo formulation^25^ of the Carreau-Yasuda viscosity model and a fluid density of 1056 *kg/m*^3^. The estimated^26,27^ peak Reynolds number of 11646 greatly exceeded the critical^26^ Reynolds number of 6959, thus the k-*ω* Shear Stress Transport (SST) Reynolds-averaged turbulence model was deployed using a low turbulence intensity (1%) at the inlet and all outlets^28^.

### 2.6 Flow Analysis

#### 2.6.1 Wall Shear Stress Indices

Using every fifth timestep from the final cardiac cycle, the Time Averaged Wall Shear Stress (TAWSS), Oscillatory Shear Index (OSI), Relative Residence Time (RRT), and Endothelial Cell Activation Potential (ECAP) were computed as follows:

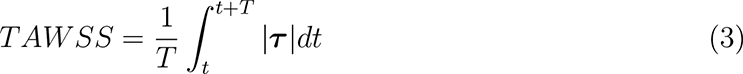

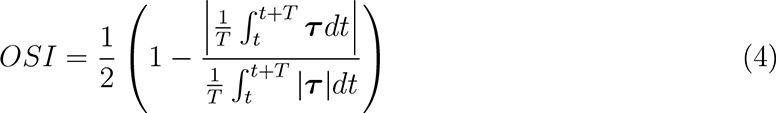

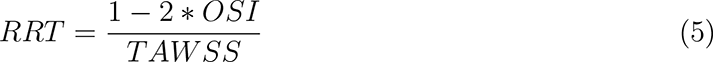

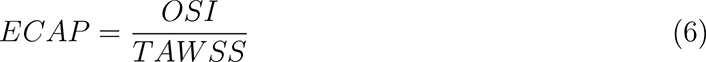

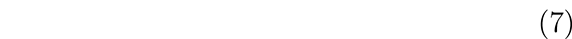

where ***τ*** (**x***, t*) is the instantaneous WSS vector and *T* is the cardiac cycle period ^29^.

#### 2.6.2 Pressure Metrics

Alongside systolic, diastolic and pulse pressures, mean transmural pressure (TMP) will be analysed in each simulation: *TMP* = *P_T L_ − P_F L_* where TL and FL pressures are evaluated as the mean instantaneous static pressure across a cross-sectional plane through the aorta. TMP magnitudes greater than 5 mmHg are associated with aortic growth in TBAD^10^.

The simulated FL ejection fraction (FLEF), calculated as the net retrograde flow volume through the PET as a proportion of the stroke volume, will also be assessed. FLEF has been associated with increased aortic growth rate in TBAD^30^.

#### 2.6.3 Helicity Metrics

We will assess the impact of IVP on bulk flow structure via qualitative and quantitative comparison of helicity. Helicity, *H*(*t*), is a scalar property used to identify streamwise vortical structures by quantifying the local alignment of velocity and vorticity vectors, **v**(**x***, t*) and ***ω***(**x***, t*), over a volume *V*:

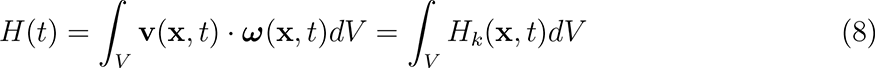

where *H_k_* is the helicity density^31^. *H*(*t*) can also be evaluated over a 2D plane by integrating the same quantities over its area. Helical flow is a natural feature of healthy aortic flow^32^ and has been demonstrated to suppress flow disturbances in cerebral aneurysms^31^. As helical structures in the aorta are on a larger scale than the boundary layer, these flow features can be measured more reliably with 4DMR than WSS measurements. As a result, associations between helicity and WSS are sought due to the potential predictive power and clinical value of WSS^31,32^.

The sign of *H_k_* indicates the direction of rotation relative to the velocity vector: positive values indicate right-handed helices (clockwise), while negative values are left-handed (anticlockwise). As *H*(*t*) = 0 can indicate either the presence of symmetrical counter-rotating vortices or zero velocity/vorticity, the magnitude of helicity can be used to distinguish these scenarios:

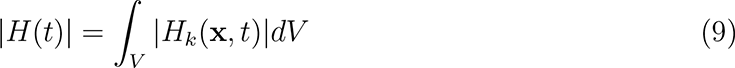

Local Normalised Helicity (LNH), defined as:

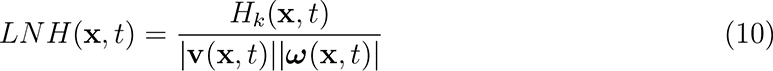

is often used to visualise vortical structures in the aorta by plotting isosurfaces of *LNH* at equal but opposing-sign values. Quantitative assessment of helicity can be performed by averaging *H_k_* over a defined volumetric region, *V*, and time interval, *T*:

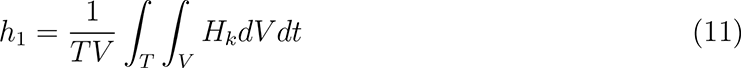

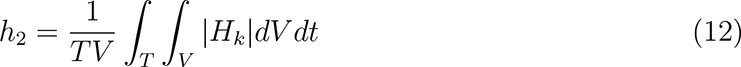

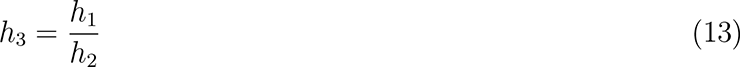

where *h*_1_ = 0 with reflectional symmetry in helical structures, or with zero velocity or vorticity. Index *h*_2_ quantifies the total amount of helicity in the volume regardless of direction. The value of *h*_3_ reflects the relative balance between right- and left-handed helicity, and its direction. Helicity indices *h*_1_-*h*_3_ were computed across the full cardiac cycle, and across systole and diastole in the five aortic subdomains depicted in Fig. 1: *AA*, *TL_t_*, *FL_t_*, *TL_a_* and *FL_a_*.

#### 2.6.4 Flow Decomposition

POD decomposes a chosen flow into a set of spatial modes, each of which is modulated by a time coefficient. In this study, we consider the flow velocity **v**(*x, y, z, t*). The fluctuating velocity about the mean is defined as 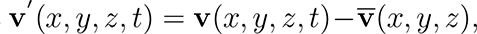 where 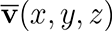 is the temporal mean of **v**(*x, y, z, t*). Applying POD, **v’** (*x, y, z, t*) is decomposed into *k* spatial modes, Φ(*x, y, z*), each associated with a time coefficient *a*(*t*), as follows:

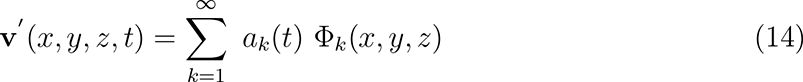

Modes are ordered by KE content. The ‘zeroth’ POD mode represents the time-averaged flow while all higher modes represent time-dependent flow structures^18^. The spatiotemporal resolution of 4DMR data is insufficient for POD analysis, so higher-fidelity 2D or 3D velocity data from Particle Image Velocimetry (PIV) or CFD are typically used in this process^18^. In this work, POD analysis was performed in MATLAB using 127 velocity snapshots throughout the full aorta at 5ms increments across the final cycle of each CFD simulation.

The energy-based ranking of modes is performed such that the flow can be accurately reconstructed using a reduced number of these modes^17^, thus reducing the order of the system. The flow is said to be accurately represented by the ensemble of modes containing 98% of the total KE which, in cardiovascular flows, has been widely reported to occur within the first 1-10 modes^16,19,33^.

Due to the energy-based nature of this analysis, modes may not retain physical interpretability^17^. Nevertheless, POD analysis offers a means of examining how the fluid KE is distributed amongst coherent flow structures and how this energy changes over time, facilitating a decoupling of spatial and temporal behaviour which can provide valuable insights into the behaviour of the flow^16^. For example, POD analysis of the flow in ruptured and unruptured aneurysms showed that rupture was more strongly associated with spatial flow complexity, rather than temporal stability^19^; similar insights may be available in the context of TBAD.

## 3 Results

In this section, time instants *T*_1_ to *T*_5_ are selected across the cardiac cycle to illustrate our key finding. These time instants are depicted schematically in Fig. 1 and correspond to mid-systolic acceleration, peak systole, mid-deceleration, early- and mid-diastole, respectively. Emphasis will be placed on the thoracic aorta as this is where aortic growth is primarily observed.

### 3.1 Velocity Distribution

Velocity magnitude contours at mid-deceleration (*T*_3_) from 4DMR and each simulation are shown in Fig. 2 and for other time instants in Supplementary Material SM2. As shown in these figures, regions with high and low velocity correspond closely between CFD and 4DMR throughout the cardiac cycle. However, velocity magnitude is universally lower in 4DMR than CFD in both lumens. 4DMR is known to perform poorly in low-velocity regions such as the FL, and to underestimate peak velocity ^9,15^. Additionally, the cycle-averaging inherent in 4DMR data may affect velocity magnitude measurements in regions experiencing highly unsteady flow (e.g. beyond the PET), and the low spatial resolution of 4DMR cannot resolve small-scale flow features. Imaging uncertainties are likely to account for many of the differences observed between 4DMR and CFD. As such, the 3D case will be considered as the baseline case herein, with each additional IVP case compared against it. This approach has been employed in other studies ^11^.

**Figure 2:**
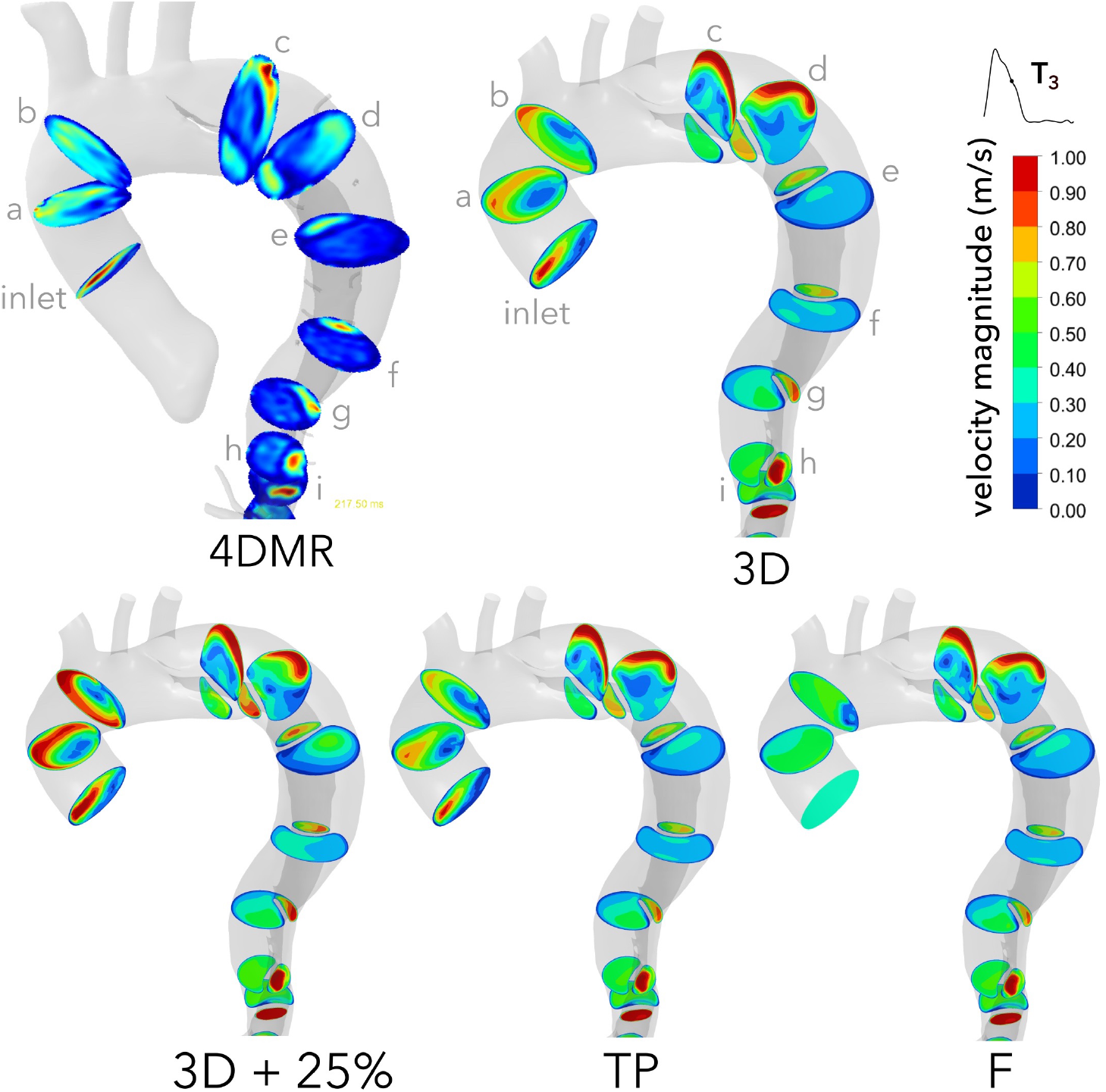
Contours of velocity magnitude in the thoracic aorta at mid-deceleration (T_3_) in each case, compared with 4DMR. Note that velocity contours are clipped to 1 m/s for clarity, but velocity reaches up to 1.3 m/s. Similar contours are provided at T_2_ and T_4_ in Supplementary Material SM2

Between cases, TL velocity distributions are qualitatively similar but with higher velocity magnitudes in the +25% case as expected. Qualitative differences between cases are primarily observed in the ascending aorta (planes *a* and *b*) and mid-thoracic FL (planes *d*-*g*). To quantify these differences, we compared the Pearson correlation coefficients of velocity magnitude on each analysis plane between each IVP and the 3D case at five points across the cardiac cycle. These results are shown in Fig. 3 and details on our calculation methods can be found in Supplementary Material SM3.

**Figure 3:**
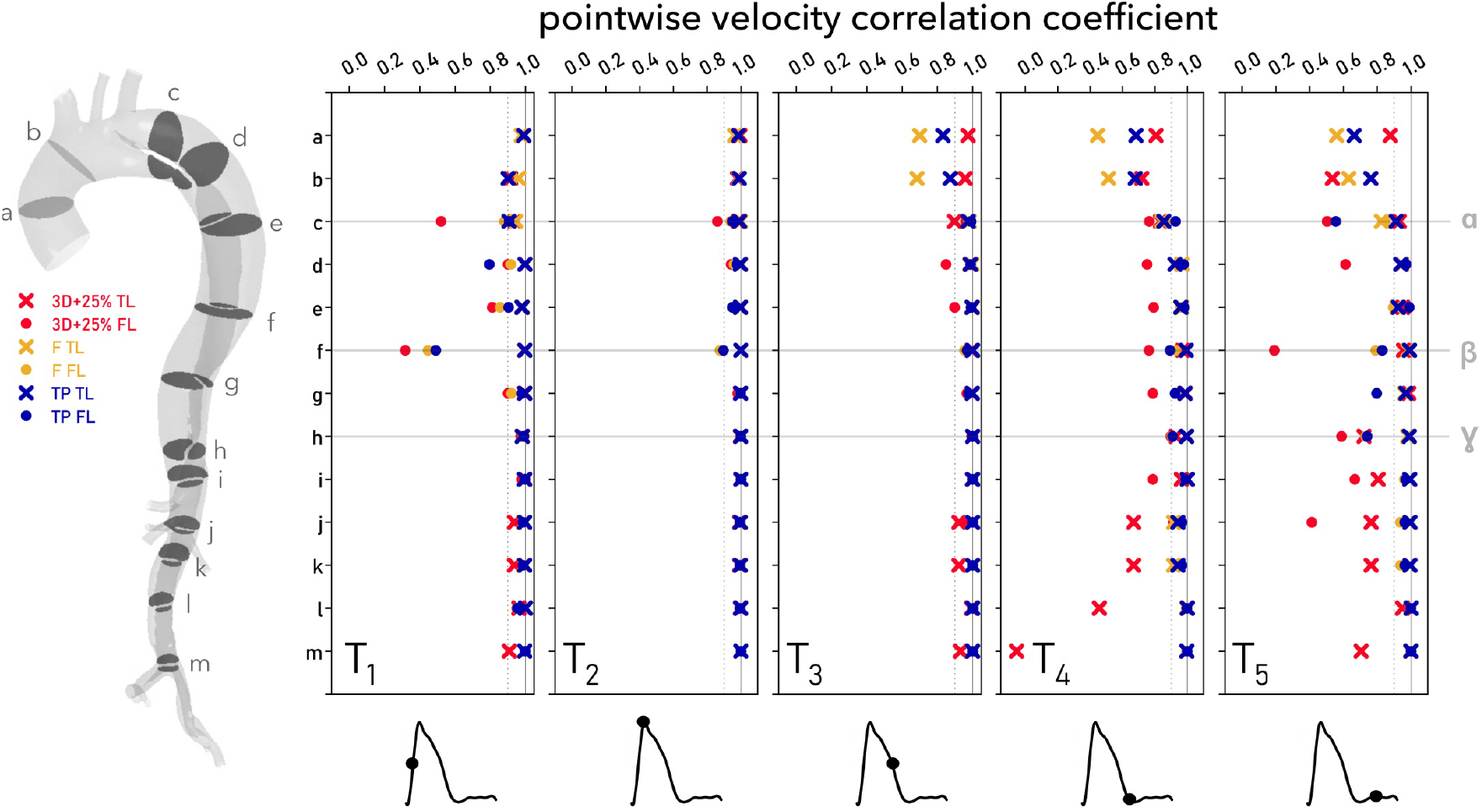
Pearson correlation coefficients between 3D IVP velocity magnitude and each other case at T_1_ to T_5_. Coefficients were computed by comparing every node on each indicated plane along the length of the dissection following the method described in Supplementary Material SM3. TL and FL points are distinguished by using different symbols, as indicated at the far left. Points α and β are the most rapidly growing regions of the FL and γ is the least, as shown in Fig. 6.

During flow acceleration (*T*_1_), velocities are well-correlated with exception to the proximal FL at plane *f*, and on plane *c* in the +25% case. At peak systole (*T*_2_), velocities are highly correlated throughout, with marginal decorrelation on planes *c* and *f*. During flow deceleration (*T*_3_), the poorest correlation is observed in the ascending aorta, with case F decorrelating most, followed by TP and +25%. During diastole, at *T*_4_ and *T*_5_, poor correlation prevails in the ascending aorta in all cases. The +25% case decorrelates particularly strongly in the FL and the abdominal TL, even exhibiting a negative correlation at *T*_4_. Throughout the cycle, planes *c* and *f*, lying within the most aneurysmal region, exhibit poor agreement in each case.

### 3.2 Pressure Metrics

Simulated pressures, TMP and FLEF are shown alongside target values in Table 2. Despite substantial aortic growth in this patient, measured FLEF is only 2.1% and mean TMP reaches only 2 mmHg.

**Table 2:** Pressure metrics from each case alongside the target measured values. † indicates that the systolic pressure measurement is derived from the brachial pressure rather than measured directly.

|  | 3D | Flat | TP | + 25% | Measurement |
| --- | --- | --- | --- | --- | --- |
| $P_s$ (mmHg) | 127.8 | 129.6 | 128.8 | 133.2 | 127<sup>†</sup> |
| $P_d$ (mmHg) | 80.4 | 80.8 | 80.9 | 83.5 | 81 |
| $P_{pulse}$ | 47.4 | 48.8 | 47.8 | 49.7 | 46 |
| FLEF (%) | 0.14 | 0.00 | 0.00 | 0.00 | 2.13 |
| Min. TMP (mmHg) | -1.93 | -1.97 | -1.96 | -2.48 |  |
| Max. TMP (mmHg) | 1.56 | 1.39 | 1.44 | 1.84 |  |

Despite identical inlet flow rate waveforms and outlet boundary conditions, F exhibits a 3% higher pulse pressure than 3D while TP exceeds 3D by less than 1%. Maximal TMP is lower with 3D than F and TP, while minimum TMP reaches higher magnitudes. This indicates that the FL is marginally more pressurised in F and TP than in 3D. TMP reaches greater magnitudes in +25 where minimum TMP is 28% lower than 3D and maximum TMP is only 18% higher. This negative shift again indicates more relative pressurisation of the FL.

### 3.3 Helicity Metrics

LNH isosurfaces during mid-deceleration (*T*_3_) are shown in the TL and FL in Fig. 4 alongside plots of *H*(*t*) and |*H*(*t*)| across the cardiac cycle. Bulk helicity indices *h*_1_ - *h*_3_ in each subdomain are provided in Table 3.

**Figure 4:**
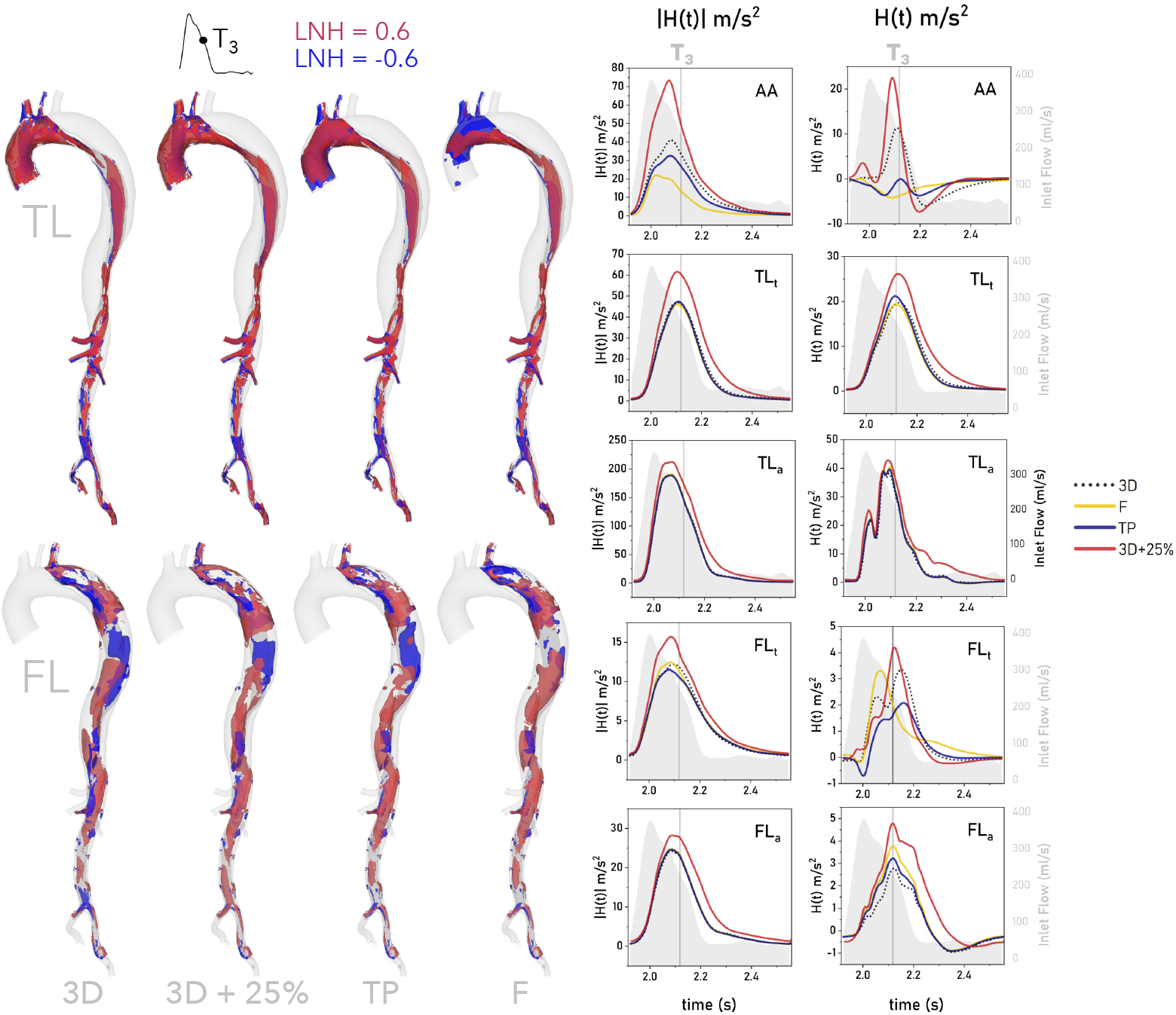
Isosurfaces of LNH = ±0.8 from each IVP case at mid-deceleration (T_3_) within the TL and FL individually (left). LNH contours at additional time points, and the time-averaged LNH, are provided in Supplementary Material SM4. Time evolution of H(t) and |H(t)| within each subdomain shown in Fig. 1 are provided on the right, with T_3_ indicated as a vertical reference line. The inlet flow rate is shown as a grey shaded region in each helicity plot using a secondary y-axis, scaled identically across all plots and shown with grey axes in all but one plot for clarity.

**Table 3:** Bulk helicity indicators h_1_-h_3_ for each IVP case. Indices are computed across systole, diastole, and the full cycle within each aortic subdomain as indicated in Fig. 1.

|  |  | systole |  |  | diastole |  |  | full cycle |  |  |
| --- | --- | --- | --- | --- | --- | --- | --- | --- | --- | --- |
| | | $h_1$ | $h_2$ | $h_3$ | $h_1$ | $h_2$ | $h_3$ | $h_1$ | $h_2$ | $h_3$ |
| **AA** | 3D | 1.61 | 23.54 | 0.07 | -2.06 | 3.67 | -0.56 | -0.22 | 13.44 | -0.02 |
|  | F | -2.21 | 11.58 | -0.19 | -0.72 | 1.11 | -0.65 | -1.43 | 6.25 | -0.23 |
|  | TP | -2.00 | 19.05 | -0.11 | -0.72 | 2.45 | -0.29 | -1.35 | 10.61 | -0.13 |
|  | +25% | 3.21 | 38.15 | 0.08 | -0.93 | 4.70 | -0.20 | 1.12 | 21.29 | 0.05 |
| **TL<sub>t</sub>** | 3D | 10.72 | 26.12 | 0.41 | 1.65 | 3.13 | 0.53 | 6.15 | 14.53 | 0.42 |
|  | F | 10.36 | 24.84 | 0.42 | 1.18 | 2.51 | 0.47 | 5.74 | 13.59 | 0.42 |
|  | TP | 10.99 | 25.36 | 0.43 | 1.28 | 2.52 | 0.51 | 6.05 | 13.75 | 0.44 |
|  | +25% | 14.04 | 34.52 | 0.41 | 3.08 | 5.94 | 0.52 | 8.46 | 19.98 | 0.42 |
| **FL<sub>t</sub>** | 3D | 1.58 | 7.73 | 0.20 | 0.01 | 1.59 | 0.01 | 0.79 | 4.63 | 0.17 |
|  | F | 1.26 | 7.65 | 0.17 | 0.29 | 1.77 | 0.16 | 0.77 | 4.69 | 0.16 |
|  | TP | 0.87 | 7.39 | 0.12 | 0.09 | 1.68 | 0.05 | 0.47 | 4.48 | 0.11 |
|  | +25% | 1.56 | 9.69 | 0.16 | -0.11 | 2.20 | -0.05 | 0.71 | 5.85 | 0.12 |
| **TL<sub>a</sub>** | 3D | 16.34 | 98.91 | 0.17 | 0.84 | 6.16 | 0.14 | 8.53 | 52.17 | 0.16 |
|  | F | 17.20 | 99.29 | 0.17 | 0.60 | 5.77 | 0.10 | 8.83 | 52.16 | 0.17 |
|  | TP | 16.84 | 97.88 | 0.17 | 0.67 | 5.74 | 0.12 | 8.62 | 51.06 | 0.17 |
|  | +25% | 19.99 | 120.17 | 0.17 | 3.92 | 12.72 | 0.31 | 11.89 | 66.02 | 0.18 |
| **FL<sub>a</sub>** | 3D | 1.18 | 13.22 | 0.09 | -0.51 | 2.06 | -0.25 | 0.33 | 7.60 | 0.04 |
|  | F | 1.73 | 13.64 | 0.13 | -0.44 | 2.01 | -0.22 | 0.63 | 7.78 | 0.08 |
|  | TP | 1.49 | 13.41 | 0.11 | -0.49 | 1.97 | -0.25 | 0.49 | 7.59 | 0.06 |
|  | +25% | 2.34 | 17.01 | 0.14 | -0.06 | 3.46 | -0.02 | 1.11 | 10.06 | 0.11 |

Helical strength, shown as a distribution of |*H*(*t*)| in Fig. 4 and characterised by the magnitude of *h*_2_ in Table 3, is greatest during the decelerating portion of the cardiac cycle^32^. Helical strength in cases 3D, F and TP match closely everywhere except the AA, where helicity is weaker when in-plane velocity components are neglected, as observed in other studies^14,34^. The +25% case provides greater helical strength throughout the domain and particularly in the AA.

Healthy aortae typically exhibit positive helicity in the AA and negative helicity beyond the mid-thoracic descending aorta ^32^. Removing in-plane inlet velocity components results in predominantly *negative* helicity in the AA during systole, evidenced by larger blue LNH volumes in Fig. 4 and negative values of *h*_1_. In contrast with healthy aortae, each descending aortic subdomain in this patient exhibit dominantly *positive* helicity (*h*_1_) throughout the cycle, except for *FL_a_* where a uniformly negative dominance develops during diastole; the 3D case presents a unique region of broader negative (blue) LNH near the iliac bifurcation that is not observed in other cases.

While minimal differences in *H*(*t*) and *h*_1_ are observed throughout the TL, except for the +25% case, which exhibits greater helicity magnitude, substantial differences are observed throughout the FL in all cases. As shown in Fig. 4, in *FL_t_*, case F exhibits an earlier peak in *H*(*t*) than the other cases. As helical strength (*|H*(*t*)*|* and *h*_2_) is very similar between 3D, F and TP cases in this region, this indicates differences in the development of right- and left-handed helical structures during diastole. Indeed, *h*_3_ values indicate a 16-fold increase in clockwise dominance in F compared to 3D.

### 3.4 Wall Shear Stress Indices

Distributions of TAWSS, OSI, ECAP and RRT in the thoracic aorta are shown for the 3D case in Fig. 5 alongside difference contours with each additional IVP case. The locations of greatest FL growth, sections *α* and *β*, are indicated on the 3D ECAP plot.

**Figure 5:**
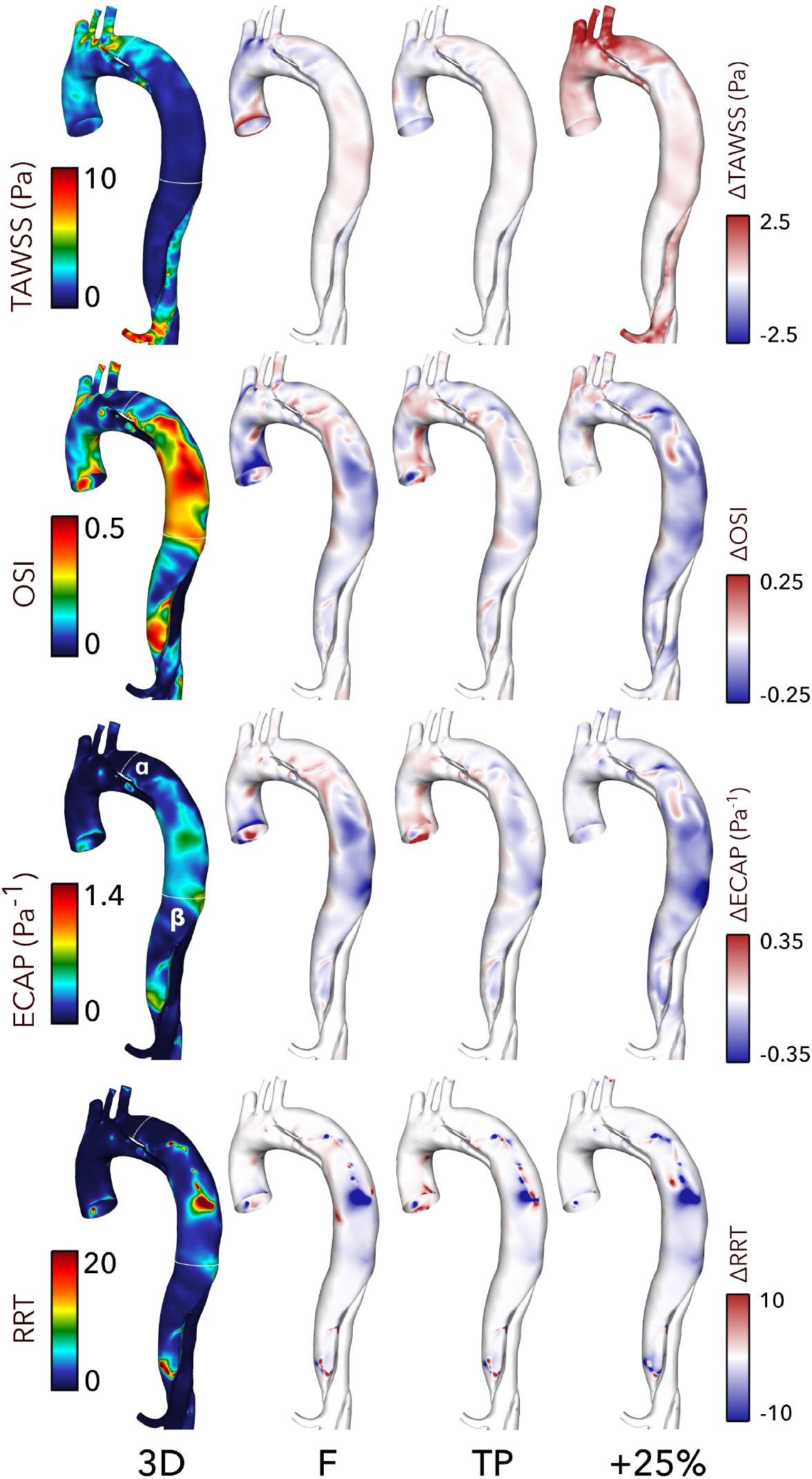
Contours of TAWSS, OSI, ECAP and RRT in the thoracic aorta from the 3D case (left) and difference contours with each other case (right). Note that contour ranges for TAWSS, ECAP and RRT are clipped for clarity of viewing. Difference contours range from 25-50% of the bounds of the 3D contours.

In the thoracic region, TAWSS is highest surrounding and immediately distal to the PET, near section *α*. TAWSS and differences in TAWSS are minimal throughout the rest of the FL. TAWSS is elevated in the +25% case, particularly in the aortic branches, due to the increased flow volume through them.

In contrast, OSI is high throughout the FL, particularly in the mid-thoracic region. While not exceeding the threshhold value of 1.4 Pa*^−^*^1^, ECAP is elevated in high OSI regions. OSI and ECAP are 20-50% lower in cases F and +25% across much of this mid-thoracic region, while TP exhibits the greatest similarity with 3D. Particularly large differences in ECAP are observed between cases at section *β*.

### 3.5 Aneurysmal Growth

We will next quantitatively compare WSS indices, time-averaged helicity 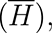 helicity magnitude 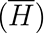 and TMP against measurements of aneurysmal growth along the thoracic FL. On cross-sectional planes co-located with FL area measurements, TAWSS, OSI, RRT and ECAP were circumferentially averaged at the wall and TMP, *H*(*t*) and *|H*(*t*)*|* were calculated as an area-average. These quantities are shown in Fig. 6, where growth is shown as a grey-shaded region. The mean discrepancy (Δ*_mean_*) and maximum discrepancy (Δ*_max_*) between each IVP and the 3D case are indicated as a percentage of the mean 3D value at each point.

**Figure 6:**
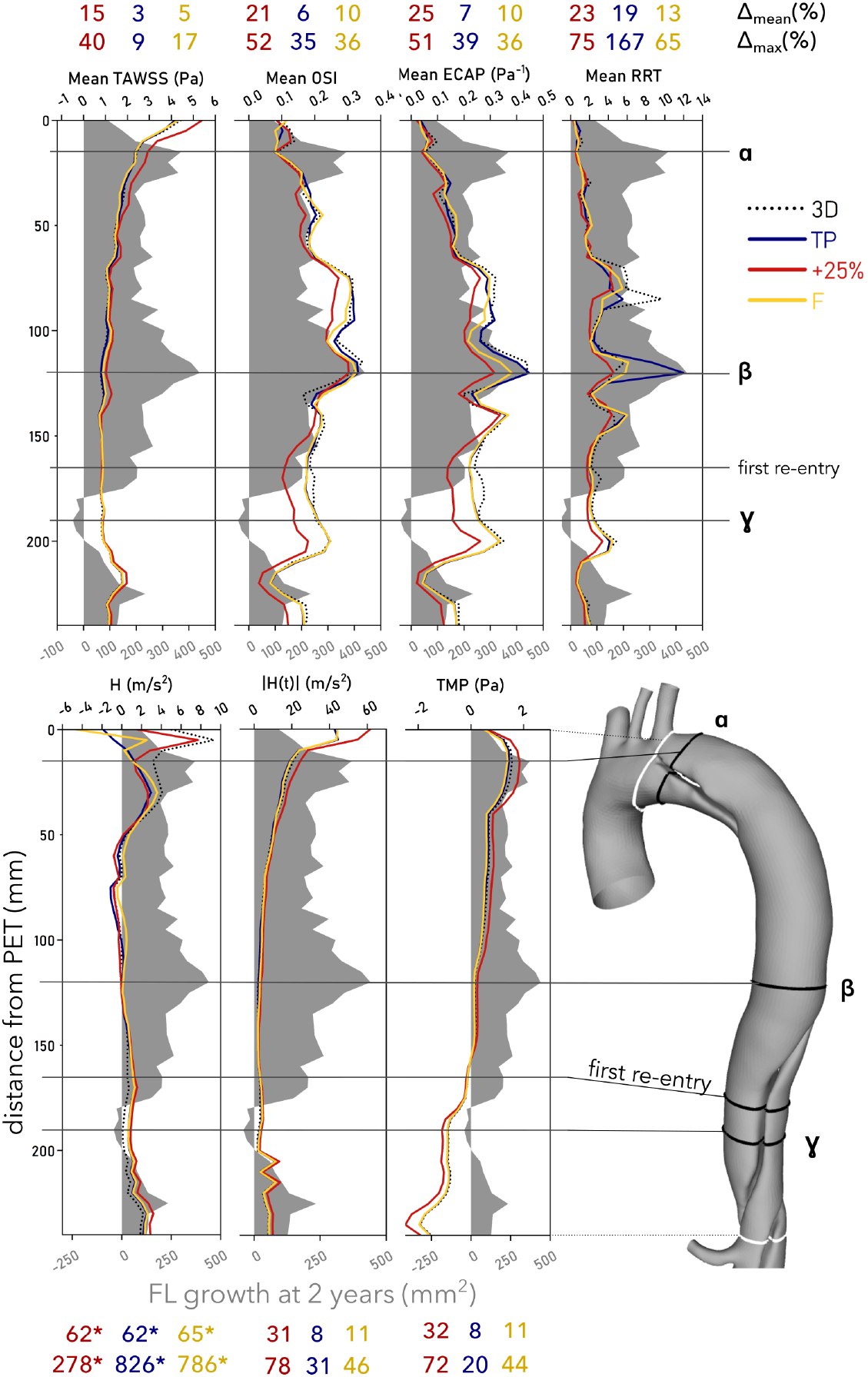
Circumferentially-averaged TAWSS, OSI, ECAP and RRT (top), plane-averaged H, |H| and TMP (bottom) are plotted along the thoracic FL as a distance from the PET against FL growth (grey shaded area) over a two-year period. Mean and maximum discrepancies (Δ_mean_ and Δ_max_) between each case and the 3D case are indicated as percentages above/below each plot. The locations of maximum growth are indicated as α and β, the location of FL regression is indicated as γ, and the location of the first re-entry tear is shown. Note that for H(t), the volumetric mean from the 3D IVP was used at each point due to the presence of many near-zero values - these of Δ_mean_ and Δ_max_ values are marked with an asterisk.

As mentioned previously, the FL grows most at cross-sections *α* and *β*, immediately beyond the PET and in the mid-thoracic FL, respectively. Marginal FL regression is observed at section *γ* in the distal thoracic FL, near the first re-entry tear.

Between IVPs, TAWSS is highest and varies most in the region immediately distal to the PET, up to *α*. Oscillatory shear is very low in this region. As TAWSS decreases along the FL, oscillatory shear increases, reaching its maximal values at section *β*, the location of greatest FL growth. Mean discrepancies are greatest in the +25% case for all WSS indices, followed by F, except RRT. Interestingly, TP exhibits greater mean and maximal discrepancies in RRT than any other case, which results from an especially high value at section *β*.

Helicity magnitude is strongest in the region proximal to the PET, reaching near-zero at *β* before increasing again beyond *γ*. Overall, the most notable differences in *H* and *|H|* between cases occur in the region proximal and immediately distal to the PET, but beyond *α* distributions are similar in all cases.

Averaged throughout the FL, TMP magnitudes are 32% higher in the +25% case than the 3D case while the flow rate-matched IVPs differ by only 8-11%. TL pressure dominates FL pressure up to the first re-entry tear, where FL pressure begins to dominate. This trend is also observed in other computational studies^35^. FL pressurisation is not directly correlated with FL growth in this patient; growing regions all exhibit different TMP characteristics; TMP is strongly positive near *α*, near-zero at *β* and negative from *γ* onward.

### 3.6 Flow Decomposition

To compare cases, the energy captured within a given POD mode was normalised by the total energy across all 127 modes within that case. Across all IVPs, the first four POD modes capture 96 *−* 97% of the total normalised KE, matching observations in other cardiovascular POD analyses^33^.

The normalised energy content of mode 1 varies between 79.4% and 84.2% for the different cases, as shown in Fig. 7, with +25% containing the least and F the most. Without normalisation, the +25% case contains more total energy in each mode than other cases across all modes due to the increased amount of KE being supplied to the system. In the second mode, the +25% case contains the most normalised energy at 12.5% while the other three contain 9.5-9.7%. In the third and fourth modes, the average normalised energy content is only around 2% and 1%, respectively, where the +25% IVP again contains more normalised energy relative to the other cases, a trend which continues across higher modes. Over 90% of the normalised energy is contained in the first two modes in all cases.

**Figure 7:**
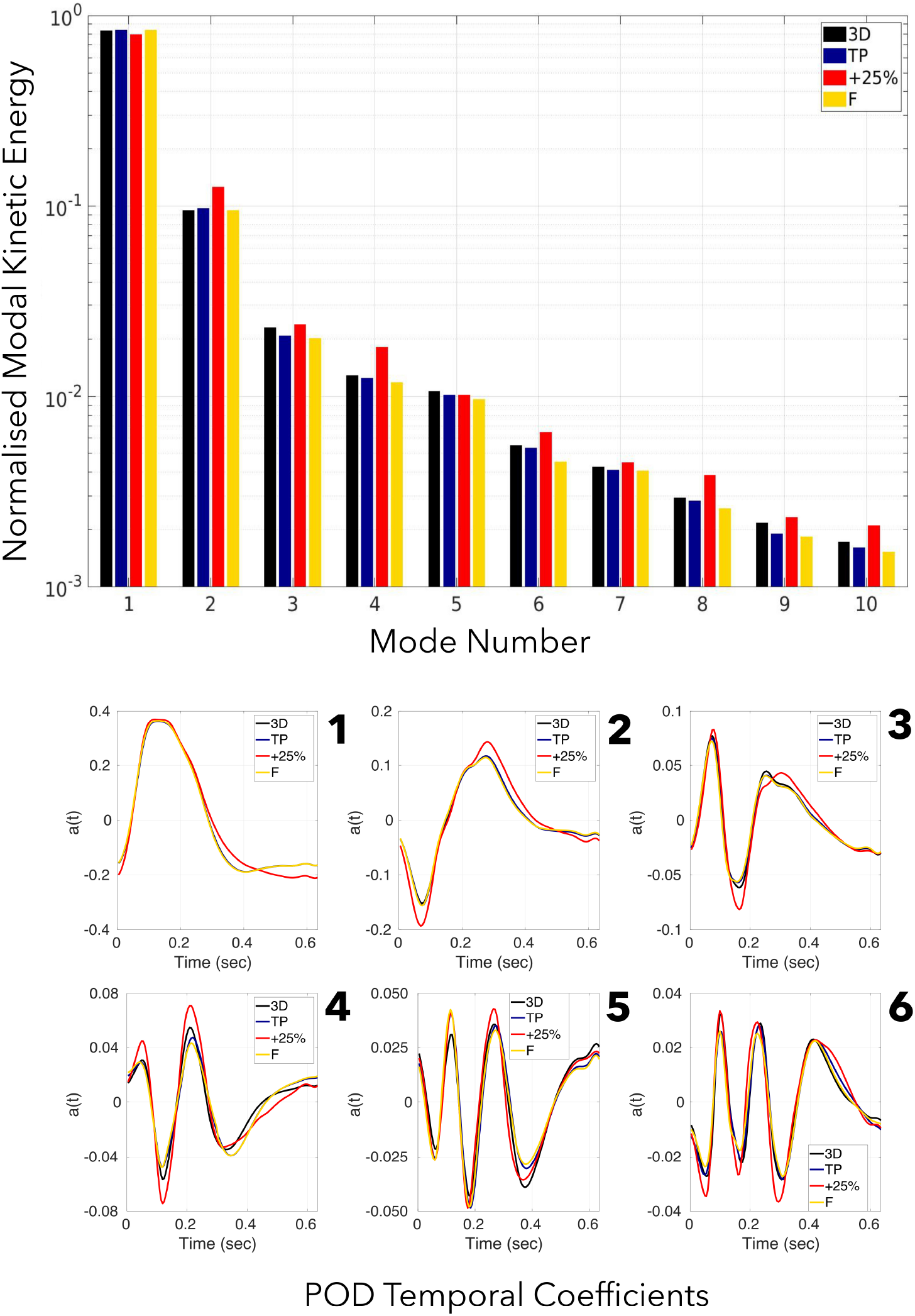
Histogram of the normalised Kinetic Energy (KE) content of the first 10 modes of the POD. Energy is normalised by the total KE across all 127 modes within each individual case.

Examining the temporal coefficients of the first six modes in Fig. 7, minimal differences are observed in flow rate-matched cases. Compared with other cases, the +25% case exhibits alterations in modes 1-4, including higher peak values and a slower decay of modes 1 and 2 during diastole.

Due to their high energy content, the impact of each IVP on the structure of modes 1 and 2 was assessed on the cross-sectional planes used previously to compare velocity magnitude contours. Contours of modes 1 and 2 are shown in Fig. 8. Similarly to the CFD velocity magnitude contours on planes a and b, shown in Fig. 2, modes 1 and 2 differ greatly in the ascending aorta across all IVPs. Differences in modes 1 and 2 become progressively less apparent along the dissection across all flow rate-matched cases with exception to planes *f* and *g* in the FL. However, the +25% case also exhibits differences across all descending aortic planes in mode 2. The 3D and TP cases exhibit the greatest similarity.

**Figure 8:**
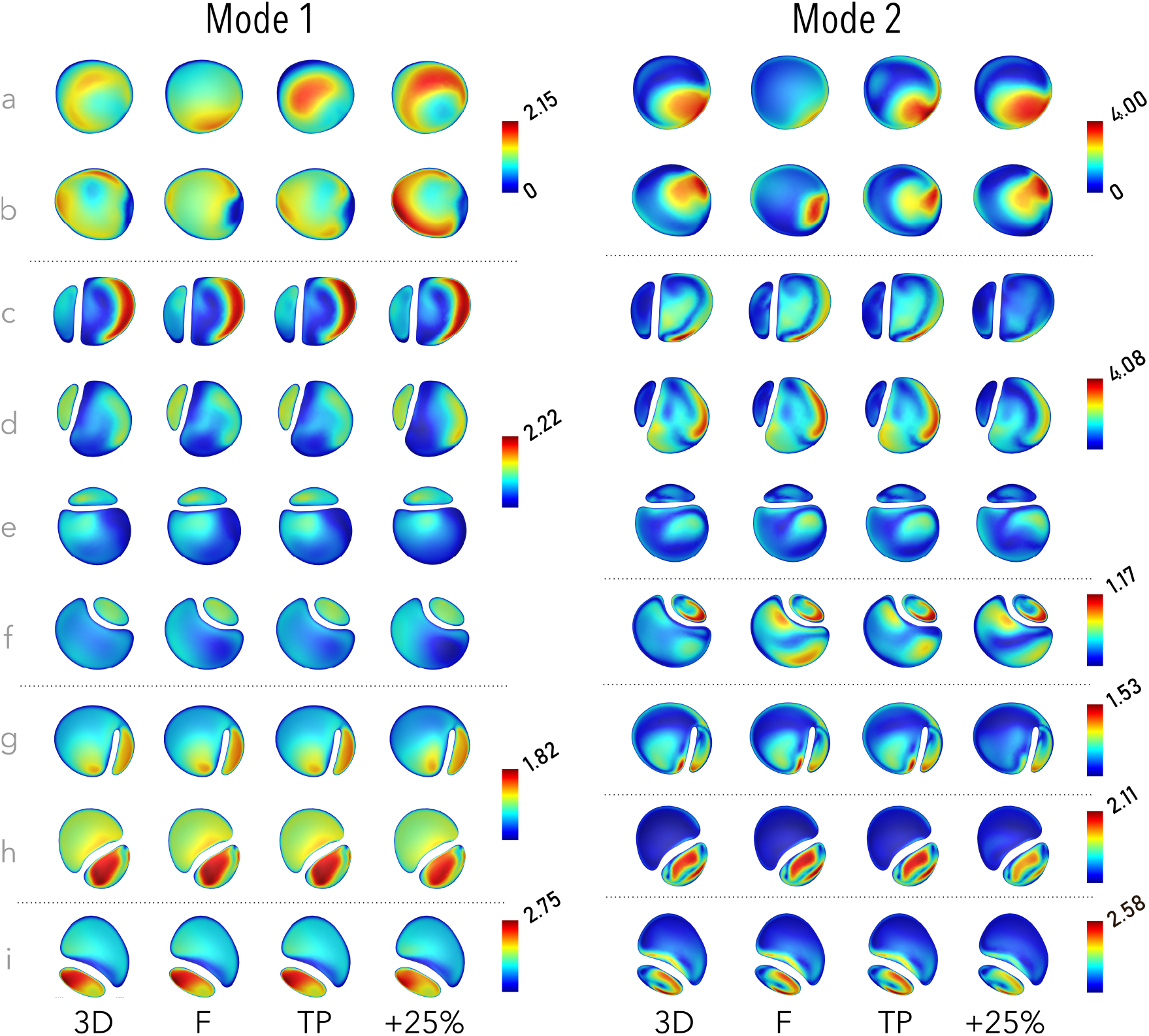
POD modes 1 and 2 on selected planes throughout the thoracic aorta (a through i, locations as shown in Fig. 2) for each of the IVP cases. Modes are a 3D vector field; these contours illustrate the magnitude of the local 3D vector. All contour scales begin at 0, shown for clarity in the top legends only. Note that planes are not to scale.

While more than 90% of the energy is contained in the first two modes, higher modes are required to accurately reconstruct the distribution of WSS. In this case, the first four modes were able to reproduce comparable distributions of TAWSS to the CFD distributions while the first seven were required to capture OSI. Reconstructed distributions of TAWSS and OSI are shown in Fig. 9, where differences between the reconstructed WSS distributions for each case can be observed to closely match the differences between CFD results with each IVP shown in Fig. 5.

**Figure 9:**
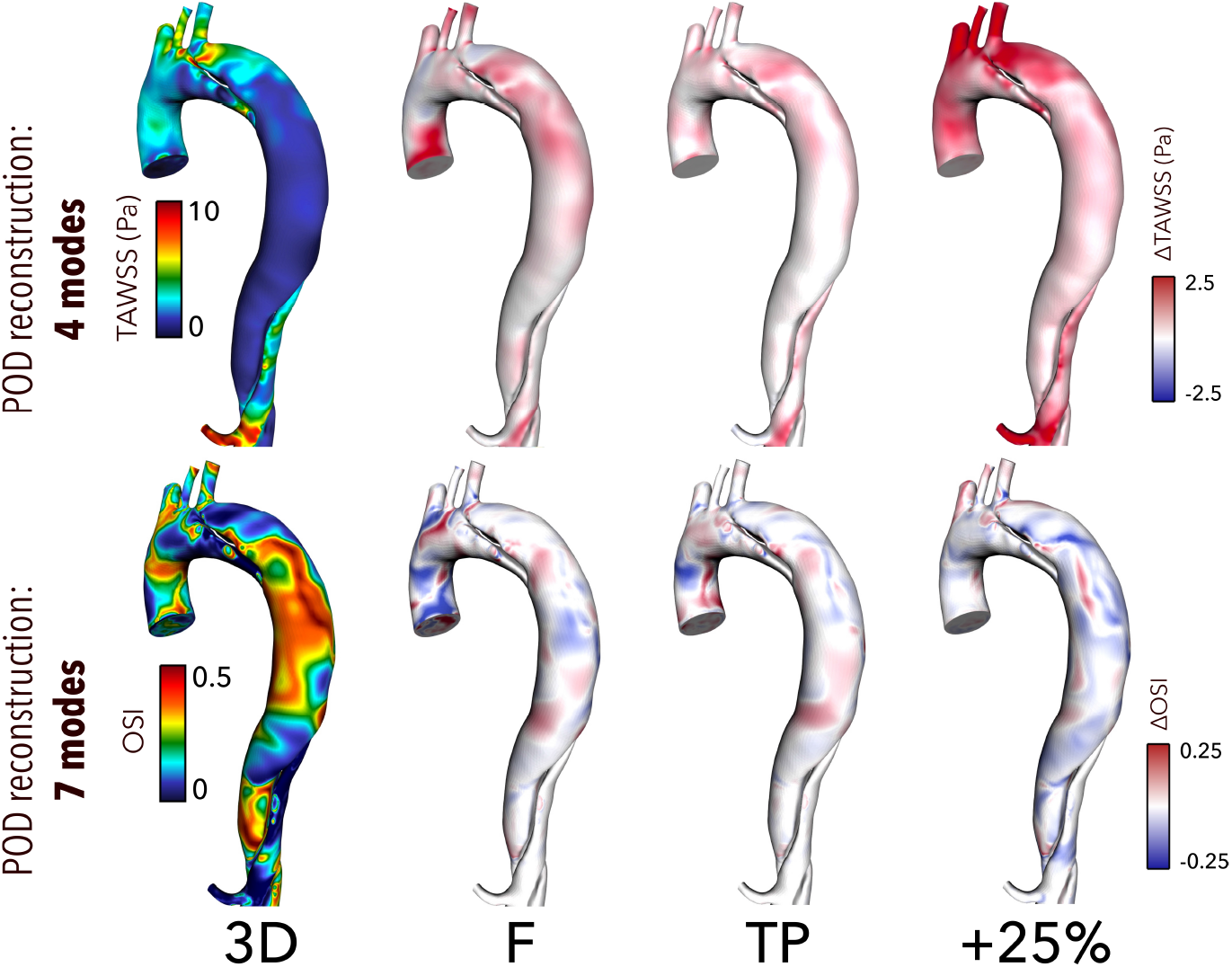
Contours of TAWSS and OSI reconstructed using the first 4 and 7 POD modes, respectively, in the thoracic aorta from the 3D case (left) and difference contours with each other case (right). Similarly to Fig. 5, note that contour ranges for TAWSS, ECAP and RRT are clipped for clarity of viewing. Difference contours range from 25-50% of the bounds of the 3D contours.

## 4 Discussion

The vast majority of patients with uncomplicated TBAD will experience aneurysmal growth and unfavourable disease progression despite best medical treatment^1^. Haemodynamic predictors extracted from 4DMR-informed CFD may assist clinicians in identifying and optimally treating these patients in future, but a more nuanced understanding of the impact of modelling assumptions, imaging errors and their combined impact on the accuracy of clinically-relevant metrics remains needed. Simulated helicity and oscillatory shear have been shown to exhibit greater sensitivity to inlet conditions than other metrics ^34,14^, but have not been examined in the context of TBAD^11,12^ despite links with disease progression^13^. In this work, we have closely examined the sensitivity of oscillatory shear and helicity to inlet conditions in a patient-specific TBAD simulation, comparing them against FL growth in this medically treated patient.

Inlet conditions are known to strongly influence velocity, helicity and WSS in the ascending aorta and arch^14,36,37,11,34^ and our results firmly corroborate these findings. However, the descending aorta is of greater interest in TBAD, where inlet conditions have been reported to affect qualitative distributions of TAWSS and velocity in the region immediately surrounding the PET whilst minimally affecting distal regions ^11,12^. With qualitative assessment alone, our results would provide similar conclusions; velocity magnitude contours and TAWSS distributions appear qualitatively similar throughout the descending aorta, regardless of IVP. However, the impact of IVP beyond the PET cannot be assessed comprehensively with qualitative analysis alone. Moreover, 4DMR cannot adequately resolve this low-velocity and highly aneurysmal region due to low measurement signal, making simulation accuracy of critical importance.

Regions of low TAWSS, which have been directly correlated with aneurysmal growth in AD^38^, are typically co-located with regions experiencing high OSI. By definition, these regions also experience elevated ECAP and RRT. We indeed observe low TAWSS and high OSI, RRT and ECAP throughout the thoracic FL. Circumferentially-averaged OSI, ECAP and RRT reach their highest levels at the location of greatest FL growth, section *β*. At this location, FL velocity is most highly decorrelated between cases during both systole and diastole. Oscillatory shear indices are therefore considerably more sensitive to IVP than TAWSS.

Being more reliably measured by 4DMR than WSS indices due to the large size of aortic helical structures relative to image resolution, flow helicity has recently been associated with FL growth in TBAD patients^13^ and has been identified as a surrogate marker of oscillatory shear in carotid arteries^31^. Helicity has also shown sensitivity to inlet conditions in healthy aortae^14^ but this effect has not been previously examined in TBAD. In the ascending aorta, we observed that helicity magnitude was reduced and stronger negative helicity developed when in-plane velocity components were neglected. In the dissected descending aorta, helicity was minimally affected by IVP in the TL but was highly sensitive to it throughout the FL; the temporal development, strength and directionality of helical structures were affected both by in-plane inlet velocity components and inlet flow volume. Time-averaged helicity and helicity magnitude reached near-zero at *β*, where oscillatory shear and aortic growth were greatest. Furthermore, in the +25% case, increased helical strength corresponded with a reduction in OSI, ECAP and RRT throughout the FL. Together, our results support previous evidence^31^ that helicity acts to stabilise flow, suggesting that this quantity may be used as a proxy for OSI in the FL using 4DMR data alone.

In contrast to section *β*, TMP, TAWSS and helicity were elevated at *α*, where considerable FL growth was also observed. In this region, flow jets through the PET and impinges on the wall. Differences in TAWSS and helicity were observed in the region surrounding section *α*, as reported in previous studies^11,12^, namely increased helicity magnitude, TAWSS and TMP in the +25% case. However, these metrics were minimally affected by the choice of flow rate-matched IVP. The contrasting haemodynamic conditions observed at at *α* and *β* support the notion that FL growth may be driven by different types of abnormal haemodynamic conditions rather than pathological values of a single metric (i.e. high OSI).

High magnitudes of TMP and high FLEF have been linked with FL growth^30,10^. Despite substantial aneurysmal growth in this patient, measured and simulated FLEF did not exceed 2.1% and simulated TMP reached only 2 mmHg, likely due to the presence of numerous large intra-luminal communications throughout the dissection which minimise pressure and flow gradients between them. Pressure-mediated vascular remodelling may be mechanistically distinct from WSS-mediated remodelling, leading to aneurysmal growth in the absence of large pressure gradients. Simulated pressure has shown high sensitivity to IVP in healthy aortae^39^. Compared with 3D, TMP was 8, 11% and 32% lower with TP, F and +25% IVPs on average, much greater than the 0.5% and 6% differences observed in a previous study^11^. The impact of IVP on pressure will be highly patient-specific, depending greatly on the size and location of luminal tears.

With analysis via POD, the +25% case exhibited the greatest differences in normalised modal energy, spatial distribution and temporal evolution. The +25% case possessed the least normalised energy in mode 1 and a relatively greater amount in higher modes compared with the flow rate-matched IVPs, indicating a higher degree of turbulence and flow complexity that has been similarly observed in other cardiovascular flows when the inlet flow rate is increased^18^. TAWSS and OSI were adequately reconstructed using only 4 and 7 POD modes in each case, capturing 96.6% and 98.7% of the normalised energy on average. Previous studies have also observed the need for higher-order modes to reconstruct OSI compared with TAWSS^40^, indicating that oscillatory shear distribution is more greatly affected by lower-energy, higher-frequency modes than TAWSS. The greater differences in higher modes in the +25% case indeed correspond to greater changes in oscillatory shear than flow rate-matched IVPs.

By reconstructing WSS distributions accurately in a reduced-order format, our work suggests that POD analysis may also offer opportunities for 4DMR data enhancement and rapid aortic flow reconstruction, providing high-fidelity haemodynamic data within clinically-relevant timescales^41,20^. Furthermore, if changes in spatial and temporal mode behaviour with increased inlet flow rate are consistent and predictable across a wider patient cohort, POD analysis may be utilised to perform efficient uncertainty quantification, perhaps in combination with machine learning techniques.

To summarise, previous studies have considered 3D IVPs from 4DMR data as the gold-standard IVP. However, upon increasing inlet velocity components by 25% to simulate the reported underestimation of velocity by 4DMR^15^, the average magnitude of differences in WSS and helicity metrics were even greater than flow rate-matched IVPs. While F captured the magnitude and trend of circumferentially-averaged WSS metrics, local values of OSI, ECAP and RRT in the most aneurysmal region (*β*) exhibited considerable differences compared with 3D, and the development of helical structures also differed greatly. Meanwhile, TP, representing an IVP derived from 2D-Flow MRI data, provided comparable results to the 3D case throughout the aorta, echoing conclusions from previous studies ^11,14^. Consequently, the greater resolution of 2D-Flow MRI and thus improved measurement of inlet flow volume may provide higher accuracy than a 3D IVP derived from 4DMR provided that the acquisition plane is appropriately chosen. Further work may endeavour to use both 2D-Flow MRI and 4DMR data to inform the IVP. Alternatively, the 4DMR imaging domain may be restricted to the ascending aorta alone to facilitate higher spatiotemporal resolution and reduce uncertainty in velocity measurements while multi-VENC 4DMR may also limit the impact of imaging errors.

The results of this study must be considered in the context of several limitations. Firstly, we have used a rigid wall assumption. Introducing aortic compliance may be expected to reduce the area exposed to high oscillatory shear and low TAWSS which are of particular interest in this study^7,42^. The rigid-wall assumption is also likely to affect simulated values of FLEF and TMP as aortic compliance acts as a pump. However, the chronicity of this case and the low measured FLEF suggest that the aorta and flap are sufficiently rigid to justify a rigid-wall assumption. Movement of the ascending aorta is also neglected, requiring a dynamic mapping of 4DMR data to the static aortic inlet which does not precisely preserve the spatial distribution of 4DMR measurements. However, the efforts taken to minimise these effects have been described and CFD velocity distributions using the 3D IVP closely match 4DMR in the ascending aorta. The artificial viscosity that results from our use of a RANS turbulence model may affect the energy distribution amongst higher POD modes, however, any such effects are likely to be minimal and should not affect our conclusions due to the vanishingly small amount of energy contained within them.

## 5 Conclusions

Our results indicate that both inlet flow volume and velocity distribution are important considerations in accurately simulating oscillatory shear and helicity throughout the FL, both of which hold predictive potential in the long-term evolution of TBAD. Patient-specific 3D IVPs extracted from 4DMR data alone may not provide sufficient accuracy due to imaging errors, despite being currently regarded as the gold-standard choice of IVP. In many cases, a TP IVP derived from 2D-Flow MRI may be more accurate. Furthermore, the highest and lowest levels oscillatory shear and helicity, respectively, were observed in the region of greatest FL growth and where IVP had the greatest impact on haemodynamic metrics.

As this study has examined only a single case of TBAD, assessing the impact of MRI-derived IVPs using CFD and POD across a wider patient cohort may provide further clarity on the best choice of IVP, illuminate novel links between coherent structures, helicity, oscillatory shear and aneurysmal growth and offer predictive metrics that may confer clinical benefit in future. As research strives toward robust, proven links between haemodynamics and disease progression, attention to patient-specific boundary conditions should be prioritised alongside the development of rapid haemodynamic simulation techniques.

## Acknowledgments

This work has been generously supported by the Department of Mechanical Engineering at University College London, the Wellcome-EPSRC Centre for Interventional Surgical Sciences (WEISS) (203145Z/16/Z), the British Heart Foundation (NH/20/1/34705), the Biotechnology and Biological Sciences Research Council (BBSRC) and UK Research and Innovation (UKRI) (BB/X005062/1), and PIONEER (EPSRC-funded, EP/W00481X/1). We also acknowledge support from the Computer Science Department at University College London and the use of their high-performance computing facilities. We finally thank Nina Montaña Brown and Yipeng Hu for their assistance with image registration.

## Notes

### Competing Interest Statement

The authors have declared no competing interest.

### Summary of Updates

Modifications to figures and restructuring of text

