## SupplementaryMaterial for "Aneurysmal Growth in Type-B Aortic Dissection: Assessing the Impact of Patient-Specific Inlet Conditions on Key Haemodynamic Indices"

#### SM1 Mesh Independence

To determine an appropriate mesh resolution for the simulations in this study, a number of relevant haemodynamic quantities were evaluated on three successively refined meshes using percentage changes and Grid Convergence Index (GCI)[SM1,SM2]. Tetrahedral meshes were generated in Fluent Mesh (ANSYS Inc., PA, USA) using sizing sensitive to proximity and curvature, a growth rate of 1.2, and maximum/minimum cell sizes as indicated in Table I. Ten near-wall layers were used in each mesh, as recommended when using the k-$\omega$ SST model. A first cell height of 0.05mm was applied in all meshes with the aim of a y+≈1 to ensure that the first cell lay within the viscous sublayer. This value resulted in a mean y+ of 0.823 across all walls, and a maximum of 3.730 at peak systole, falling well within the recommended value of 5.

GCI was calculated as a percentage using the following equations, where $c$, $m$ and $f$ correspond to quantities from the coarse, medium and fine meshes, respectively:

$$r_{f,m} = \left(\frac{N_f}{N_m}\right)^{\frac{1}{3}} \approx r_{m,c} = \left(\frac{N_f}{N_m}\right)^{\frac{1}{3}}$$

$$r = \frac{r_{f,m} + r_{m,c}}{2}$$

$$p = \frac{ln\left(\frac{|f_c - f_m|}{|f_m - f_f|}\right)}{ln(r)}$$

$$E_{f,m} = \frac{\left(\frac{|f_m - f_f|}{f_f}\right)}{r^p - 1} \qquad E_{m,c} = \frac{\left(\frac{|f_c - f_m|}{f_m}\right)}{r^p - 1}$$

$$GCI_{f,m} = F_S|E_{f,m}| \qquad GCI_{m,c} = F_S|E_{m,c}|$$

$N$ is the number of elements in the mesh, $f$ is the examined variable of interest, and $F_S$ is a safety factor of 1.25[SM1,SM2,SM3].

We compared systolic and diastolic pressure at the inlet, maximum velocity in the full volume at peak systole, area-averaged WSS magnitude on the wall at peak systole, maximum and mean OSI and TAWSS. Across all meshes, GCI did not exceed 3.1% for any quantity in any mesh. Between fine and medium meshes, the maximum difference in any quantity was

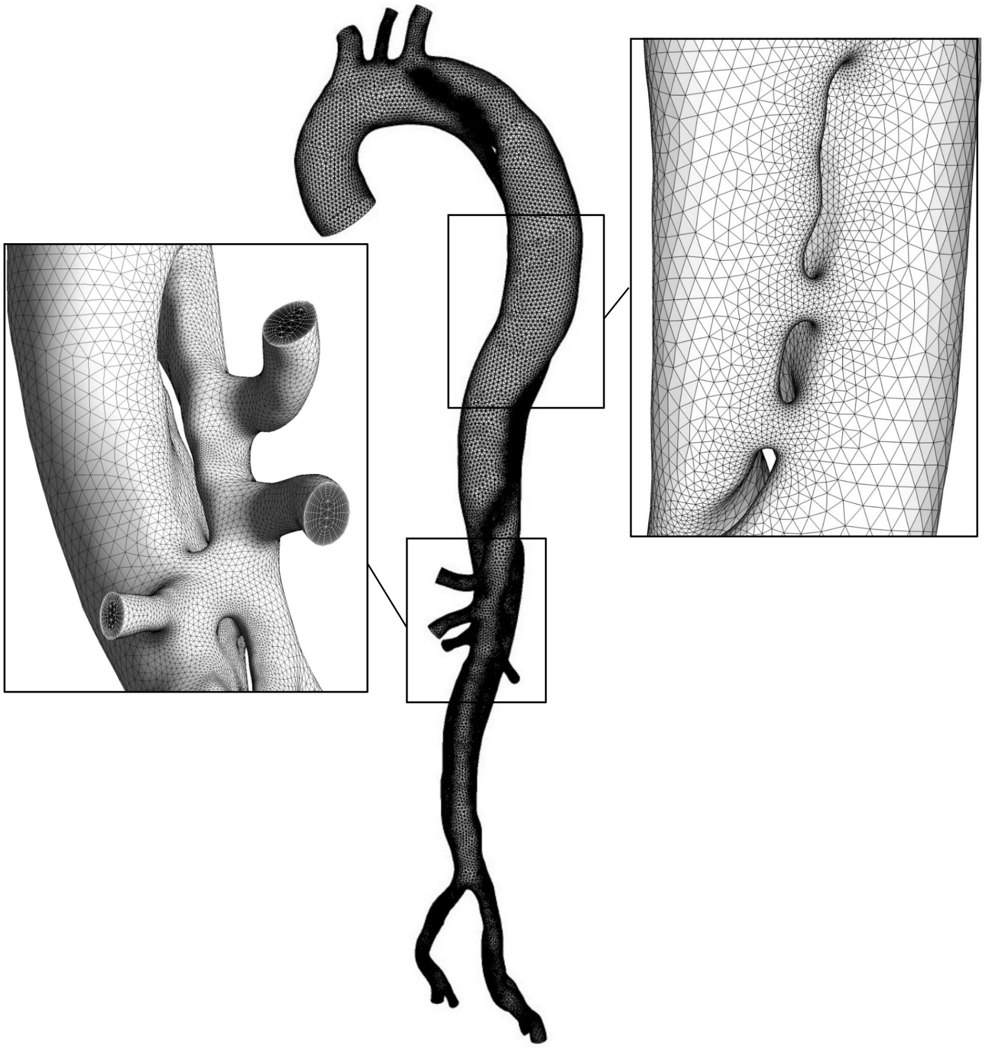

Figure I: *Images of the final mesh (medium), with detail views of the branches and tears.*

| | $f$ | $m$ | $c$ | $\%_{f,m}$ | $\%_{m,c}$ | $GCI_{f,m}$ | $GCI_{m,c}$ |
|---|---|---|---|---|---|---|---|
| $cell_{min}$ $(mm)$ | 0.4 | 0.8 | 1 | 100 | 25 | | |
| $cell_{max}$ $(mm)$ | 1 | 1.5 | 2 | 50 | 33.33 | | |
| $Elements$ | 5020766 | 2298005 | 1310323 | -54.23 | -42.98 | | |
| $P_S$ | 129.1 | 127.8 | 130.6 | -1.01 | 2.19 | **1.091** | 2.374 |
| $P_D$ | 80.7 | 80.1 | 81.2 | -0.74 | 1.37 | **1.115** | 2.06 |
| $v_{max}$ | 2.135 | 2.1168 | 2.106 | -0.85 | -0.51 | **2.629** | 1.578 |
| $\tau_{max}$ | 3.5565 | 3.6155 | 3.7559 | 1.66 | 3.88 | **1.504** | 3.519 |
| $OSI_{max}$ | 0.4985 | 0.4963 | 0.4981 | -0.44 | 0.36 | **3.034** | 2.493 |
| $OSI_{mean}$ | 0.1536 | 0.1509 | 0.1514 | -1.73 | 0.36 | **2.709** | 0.56 |
| $TAWSS_{max}$ | 16.612 | 16.3276 | 155.512 | -1.71 | 852.45 | **0.004** | 2.182 |
| $TAWSS_{mean}$ | 1.3932 | 1.3979 | 1.4183 | 0.34 | 1.46 | **0.126** | 0.546 |

Table I: *Mesh properties, haemodynamic quantities of interest, and GCI values for the coarse, medium and fine meshes.*

<2% while the maximum difference between medium and coarse meshes was 852%, so the medium mesh, shown in Fig. I, was selected for the final study.

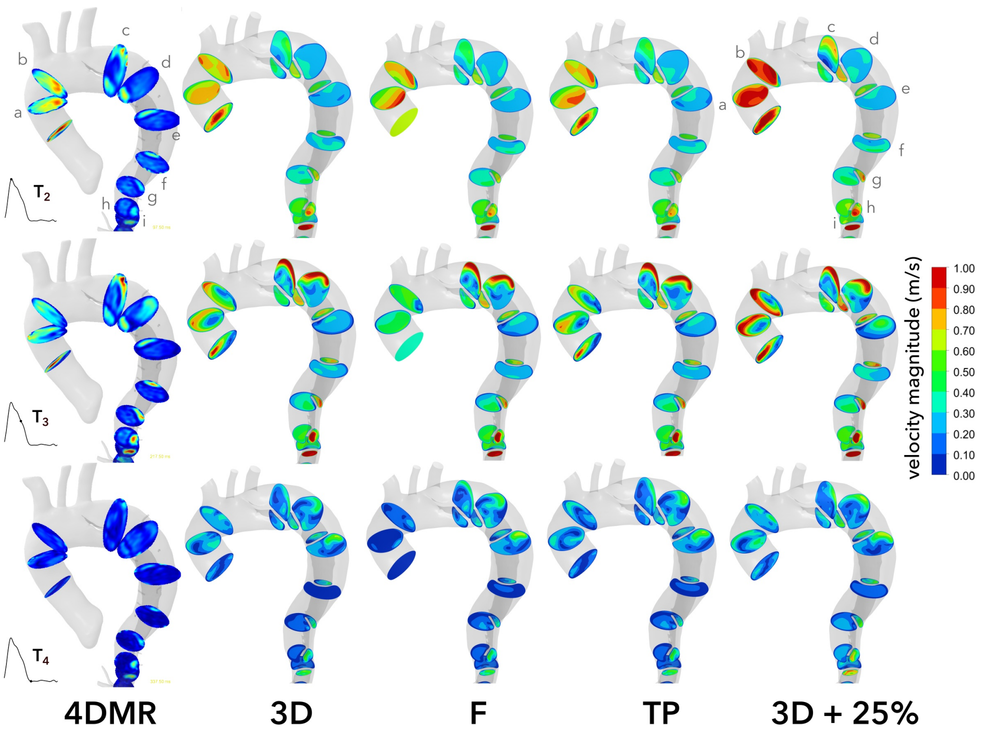


Figure II: *CFD velocity magnitude contours from each IVP case compared with 4DMR data at each time point omitted from the main text.*

### SM2 Velocity Contours

Velocity contours at $T_2$, $T_3$, and $T_4$ from 4DMR and each CFD simulation are shown in Fig. II.

### SM3 Velocity Pearson Correlation

The quantitative velocity agreement between cases was assessed using Pearson correlation coefficient. If each variable has N scalar observations, then the Pearson correlation coefficient is defined as:

$$\rho(A,B) = \frac{1}{N-1}\sum_{i=1}^{N}\left(\frac{A_i-\mu_A}{\sigma_A}\right)\left(\frac{B_i-\mu_B}{\sigma_B}\right) \tag{S1}$$

where $\mu_A$ and $\sigma_A$ are the mean and standard deviation of $A$, respectively, and similarly for $B$[SM4]. Pearson correlation provides a numerical measure of the relationship between $A$ and $B$ with values of 1 and -1 indicating a perfect positive or negative linear relationship between them and zero indicating no relationship. Note that a value of 1 does not indicate that the two independent variables are identical in magnitude for all observations, just that they follow a linear relationship.

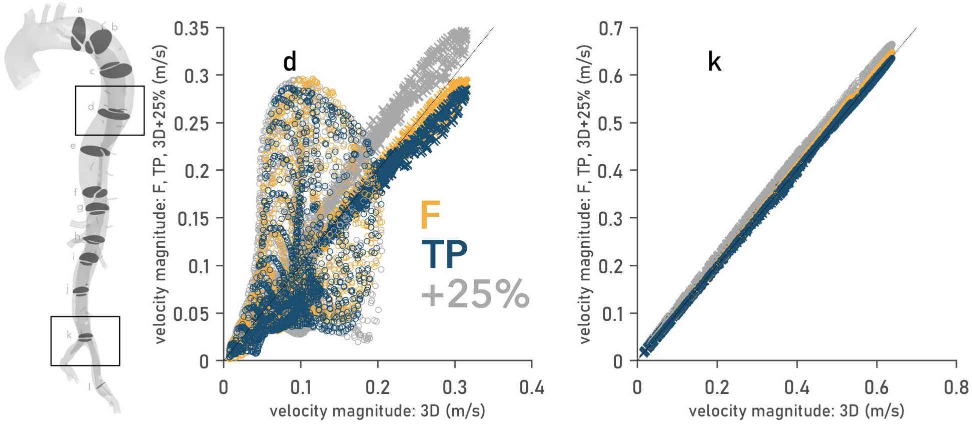


Figure III: *Pointwise velocity magnitude comparisons on planes d and k at peak systole between the 3D case and every other IVP. The x-axis represents the column matrix A described in SM3, while the y-axis represents B.*

In this case, the $A$ and $B$ are the velocity magnitudes at each point on a given plane in the 3D IVP case ($A$) and any other given IVP ($B$), at a chosen time instant. The $N$ observations are represented by corresponding points on the plane. Pearson correlation coefficients were

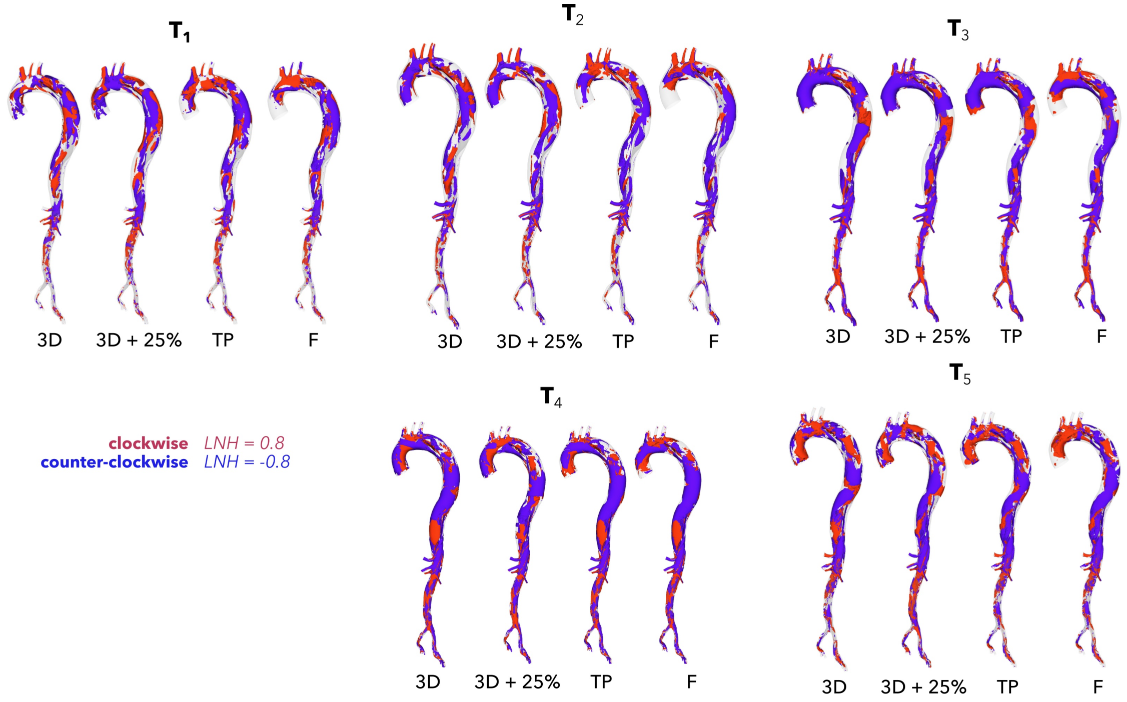


Figure IV: *LNH isosurfaces at additional time points to those provided in the main text.*

evaluated using the *corcoeff* function in MATLAB. Fig. III provides a visualisation of this data by plotting $A$ on the x-axis and $B$ on the y-axis at peak systole on planes $d$ and $k$, with a dotted line to represent $x = y$. As Pearson correlation is a measure of linearity between datasets, rather than equality, it provides an ideal quantitative measure of agreement in flow distribution when flow volumes are not precisely matched, in this case between the 3D and +25% cases.

### SM4 Helicity Isosurfaces

LNH isosurfaces are provided at additional time points in figure IV, while time-averaged LNH isosurfaces are provided in V.

[SM2] Craven, B., Paterson, E., Settles, G. & Lawson, M. Development and verification

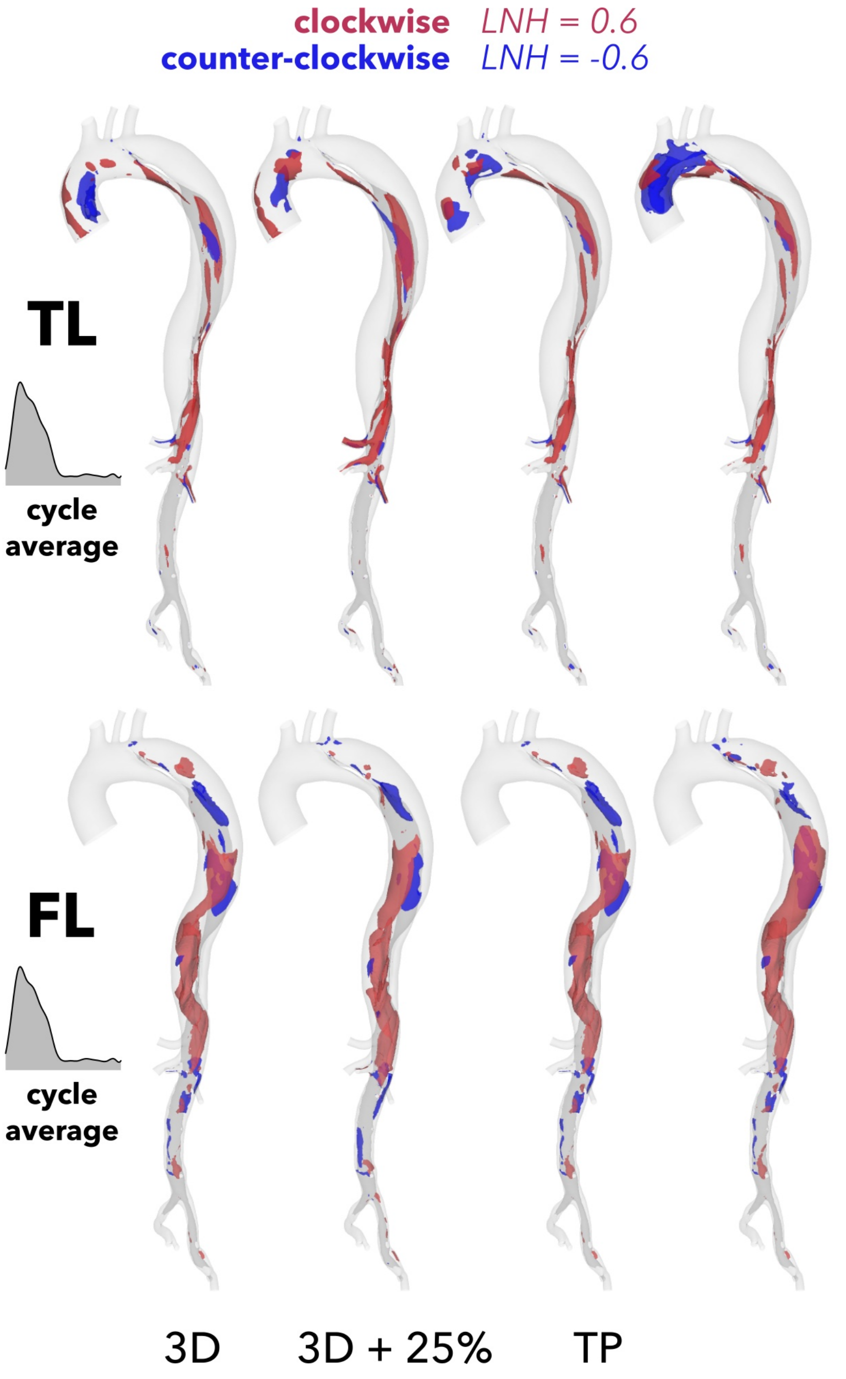


Figure V: *Mean LNH isosurfaces across the cardiac cycle.*

of a high-fidelity computational fluid dynamics model of canine nasal airflow. *J. Biomech. Eng.*. **131**, 091002 (2009)

[SM3] Armour, C., Guo, B., Pirola, S., Saitta, S., Liu, Y., Dong, Z. & Xu, X. The influence of inlet velocity profile on predicted flow in type B aortic dissection. *Biomech. Model. Mechanobiol.*. **20**, 481-490 (2021)

[SM4] MATLAB Inc., **corrcoeff**, *MATLAB R2022b Documentation*, https://www.mathworks.com/help/matlab/ref/corrcoef.htmld124e272811
